# RNAi-mediated knockdown of two orphan G-protein coupled receptors reduces fecundity in the yellow fever mosquito *Aedes aegypti*

**DOI:** 10.1101/2023.03.30.534942

**Authors:** Nia I. Keyes-Scott, Kyle R. Swade, Lena R. Allen, Kevin J. Vogel

## Abstract

G-protein coupled receptors (GPCRs) control numerous physiological processes in insects, including reproduction. While many GPCRs have known ligands, orphan GPCRs do not have identified ligands in which they bind. Advances in genomic sequencing and phylogenetics provide the ability to compare orphan receptor protein sequences to sequences of characterized GPCRs, and thus gain a better understanding of their potential function. Our study investigated two orphan GPCRs, AAEL003647 and AAEL019988, in the yellow fever mosquito, *Aedes aegypti*. From our phylogenetic analysis, we found that AAEL003647 is orthologous to the SIFamide-2/SMYamide receptor, while AAEL019988 is orthologous to the Trapped in endoderm (Tre1) receptor of *Drosophila melanogaster*. Tissue-specific expression analysis revealed that both receptors had highest expression in the ovaries, suggesting they may be important for reproduction. We then used RNA interference (RNAi) to knock down both genes and found a significant reduction in the number of eggs laid per individual female mosquito, suggesting both receptors are important for *Ae. aegypti* reproduction.

## 1. Introduction

Mosquitoes are a persistent threat to global health due to their ability to transmit pathogens among vertebrate hosts through blood feeding, which is required for many mosquito species to produce eggs. The events beginning with blood meal digestion and ultimately leading to egg production are coordinated by several reproductive hormones, including insulin-like peptide 3 (ILP3) and ovary ecdysteroidogenic hormone (OEH), which are released shortly after a blood meal is consumed^1–3^. Release of ILP3 from brain neurosecretory cells stimulates blood meal digestion, and ILP3 and OEH both stimulate secretion of 20-hydroxyecdysone (20E) from the ovaries ^1–4^. After 20E is released into the hemolymph, expression of yolk protein precursors (YPP) in the fat body is induced, initiating the production of yolk proteins, including vitellogenin, which are subsequently transported to the ovaries and packaged into oocytes resulting in egg formation^5,6^.

Hormone signaling pathways have been exploited to control insect populations. Insect chemical growth regulators (IGRs), such as 20E antagonists, target insect hormonal pathways and have been utilized to control insect disease vectors^7,8^. IGRs are attractive control measures due to their selective toxicity against insects and decreased rate of insecticide resistance developed against them relative to traditional pesticides^9,10^. IGR targets such as, JH and 20E and their receptors, are widely conserved in insects increasing the chances of negative effects on non-target species ^7,8,11–13^. An attractive alternative to IGRs that act on JH or 20E are compounds that selectively target hormones or hormone receptors that are not widely conserved across all insect groups. G-protein coupled receptors (GPCRs) and their ligands may present taxa-specific targets, as insect genomes often encode unique GPCRs, including many that bind peptide hormones that regulate important aspects of insect physiology^14–16^.

Hormone-binding GPCRs are essential in modulating insect physiology, including in metabolism^17,18^, reproduction^19^, behavior^20^, immunity^21^, and embryonic development^22^, as they transduce systemic hormonal signals into target cells. In addition to modulating a diverse number of functions in insects, GPCRs are the largest class of receptor and bind a variety of ligands, including neurotransmitters^23^ and peptide hormones^24,25^. Peptide hormones govern many physiological functions in insects including feeding^26–29^, mating behavior^30^, development^31–33^, metabolism ^1,18,34–36^, immunity^37^, diuresis^38–40^, and reproduction^1,19,41^. While the ligands of many GPCRs have been identified, genomes of even well-studied organisms still encode GPCRs whose ligands are unknown.

Comparative genomics and phylogenetic analyses are useful tools in the identification of ligands of former orphan receptors^3,19^. Phylogenetic placement of orphan receptors, such as in the case of the OEH receptor of *Aedes aegypti* mosquitoes, can provide insights into potential ligands. A Venus flytrap domain-containing receptor tyrosine kinase was found to be closely related to the mosquito insulin receptor, and also displayed the same species distribution pattern as neuroparsin peptide hormones including OEH. Subsequent biochemical and molecular studies determined that the gene in question was an OEH receptor^3^. Tissue-specific expression patterns are also useful in determining the functional roles and ligands of hormone receptors. We identified that the neuropeptide CNMa and its receptor, CNMaR, which were first identified in *Drosophila melanogaster*, were specifically expressed in *Ae. aegypti* ovaries and hypothesized that it was likely important for reproduction^3,19,42^. In Culicidae, the CNMa receptor underwent gene duplication, resulting in two receptors, CNMaR-1a and CNMaR-1b, which both actively bind CNMa in vitro^19^. In *Ae. aegypti*, CNMa and CNMaR-1b are highly expressed in female reproductive tissue and modulate the production of eggs^19,43^.

We chose to examine two orphan GPCRs of *Aedes aegypti*, AAEL003647 and AAEL019988. These orphan GPCRs were chosen for further investigation based on their expression in female reproductive tissues following a blood meal^43^, suggesting a potential role in the modulation of reproductive physiology. We built phylogenetic trees to identify closely related receptors and provide insight into possible functions of the receptors. To understand the tissue tropism and temporal distribution of AAEL003647 and AAEL019988, we conducted a detailed expression analysis of both GPCRs in juvenile and adult mosquitoes. Using RNAi, we then investigated the functional consequences of silencing the GPCRs on fecundity. These results shed new light on the role of these orphan GPCRs on the reproductive physiology of *Ae. aegypti* mosquitoes.

## 2. Materials and Methods

### 2.1 Mosquitoes

UGAL strain *Aedes aegypti* were used for all experiments. Mosquito colonies were maintained at 27°C on a 16:8h L:D cycle. Larvae were fed Cichlid Gold fish pellets (Hikari, USA, Hayward, CA), and adult mosquitoes were fed an 8% sucrose solution until 2 days post-emergence. Adult females were fed defibrinated rabbit blood (Hemostat Laboratories, Dixon, CA, USA) by an artificial feeding apparatus warmed to 37°C.

### 2.2 Phylogenetic analysis

Putative AAEL003647 and AAEL019988 orthologs were identified using BLASTp against the National Center for Biotechnology Information non-redundant protein database limiting hits to insects (table S1). Other orthologs were identified in OrthoDB^44^. Select orthologs of AAEL003647 identified by Veenstra (2021) were also included. Protein sequences were aligned using MAFFT version 7.450 software in Geneious 11.1.4^45,46^. Gaps in alignments were manually removed, and trimmed alignments were used to construct maximum likelihood phylogenies using the default parameters of PhyML^47^. FigTree version 1.4.4 was used for visualization of trees.

### 2.3 Expression profiles

Eight to ten-day old, non-blood fed mated females were collected and dissected into head, gut, fat body and abdominal carcass (“pelt”), and ovaries in sterile, nuclease-free, *Aedes* saline. Additional ovary samples were collected from females at 2-hour intervals post-feeding (pbf) until 12 hours, then at 24, 48, and 72 hours pbf. Four or more tissue samples were collected for each tissue and time point. After collection, tissue samples were stored at −80°C prior to RNA extraction. Tissue samples were thawed on ice and homogenized with a rotor pestle. Total RNA was isolated from homogenized tissues using the RNeasy Mini kit (Qiagen, Venlo, The Netherlands) according to manufacturer instructions. DNA was removed from each RNA sample using the Turbo DNA-free kit (Ambion, Austin, TX, USA). One hundred nanograms of RNA was used as input to synthesize cDNA using the iScript cDNA synthesis kit (BioRad, Hercules, CA, USA). cDNA templates were used for quantitative real-time PCR, with the Quantifast SYBR Green PCR kit (Qiagen) and gene specific primers (table S2). Standard curves for each gene were generated by cloning qPCR products into the pSCA vector with the Strataclone PCR cloning kit (Agilent, Santa Clara, CA, USA), isolating plasmid DNA using the GeneJET Plasmid Miniprep Kit (Thermo Scientific, Vilnius, Lithuania), and preparing plasmid standards to a known copy number. Expression levels of ribosomal protein S7 were used as a housekeeping gene to normalize transcript abundance.

### 2.4 RNAi knockdown of receptors and bioassays

A 400-500 bp region of each gene was chosen as a target for dsRNA synthesis for AAEL003647 and AAEL019988, subsequently referred to as *ds3647* and *ds19988*, respectively. Primers including the T7 promoter sequence were used to amplify each target using cDNA synthesized from RNA isolated from whole body, non-blood fed females (table S2). PCR products were cloned into the pSCA vector and plasmid DNA was extracted using methods listed above. Plasmid DNA from each target and an EGPF control were used as the templates for dsRNA synthesis. dsRNA was synthesized using the MEGAscript RNAi kit (Ambion, Vilnius, Lithuania), according to manufacturer instructions. Following dsRNA synthesis, dsRNA was precipitated in ethanol and resuspended in *Aedes* saline to a concentration of 2µg/µL.

Newly emerged (≤ 1d post eclosion) mated females were injected with 2 µg *ds3647, ds19988*, or *dsEGFP*. To validate receptor knockdown, whole body females were collected 7 days post-injection. qPCR was used to validate knockdown of each gene using the methods detailed above. Females were blood fed three days post-injection and separated into individual egg laying chambers consisting of a damp paper towel in a plastic cup with a lid and a dental wick with 8% sucrose solution, for yolk deposition and fecundity bioassays. For yolk deposition bioassays, females were collected at 24, 48, and 72 hours PBF. Ovaries were dissected and yolk deposition per oocyte was measured along the anterior-posterior axis using an ocular micrometer. Five oocytes were measured and averaged per female, and 5 females were used per time point and treatment. Egg laying was measured by providing females with a wet paper towel at 72 h post blood feeding to stimulate egg deposition. Females were given 48 h to deposit eggs. After 48 h hours, the number of eggs laid per individual female was counted.

## 3. Results

### 3.1 Phylogenetic comparison of AAEL003647 and AAEL019988

Our phylogenetic analysis included genomic sequences from diverse insect species to identify the closest receptor relatives across both holometabolous and hemimetabolous insects^48^, including putative SMYamide receptor sequences identified by Veenstra 2021^49^. Our results indicate a strongly supported clade of receptors that are distinct from, but sister to, the SIFamide receptors (figure 1A). These receptors are found in the genomes of culicids as well as cockroaches (*Periplaneta americana* and *Blattella germanica*), termites (*Zootermopsis nevadesis*) brown rice planthoppers (*Nilaparvata lugens)*, and *Bombyx mori* (figure 1A). This robustly supported sister clade to SIFamide receptors suggest an ancient split between SIFamide receptors and the orthologs of AAEL003647 which predates the split of hemi- and holometabolous insects.

**Figure 1.**
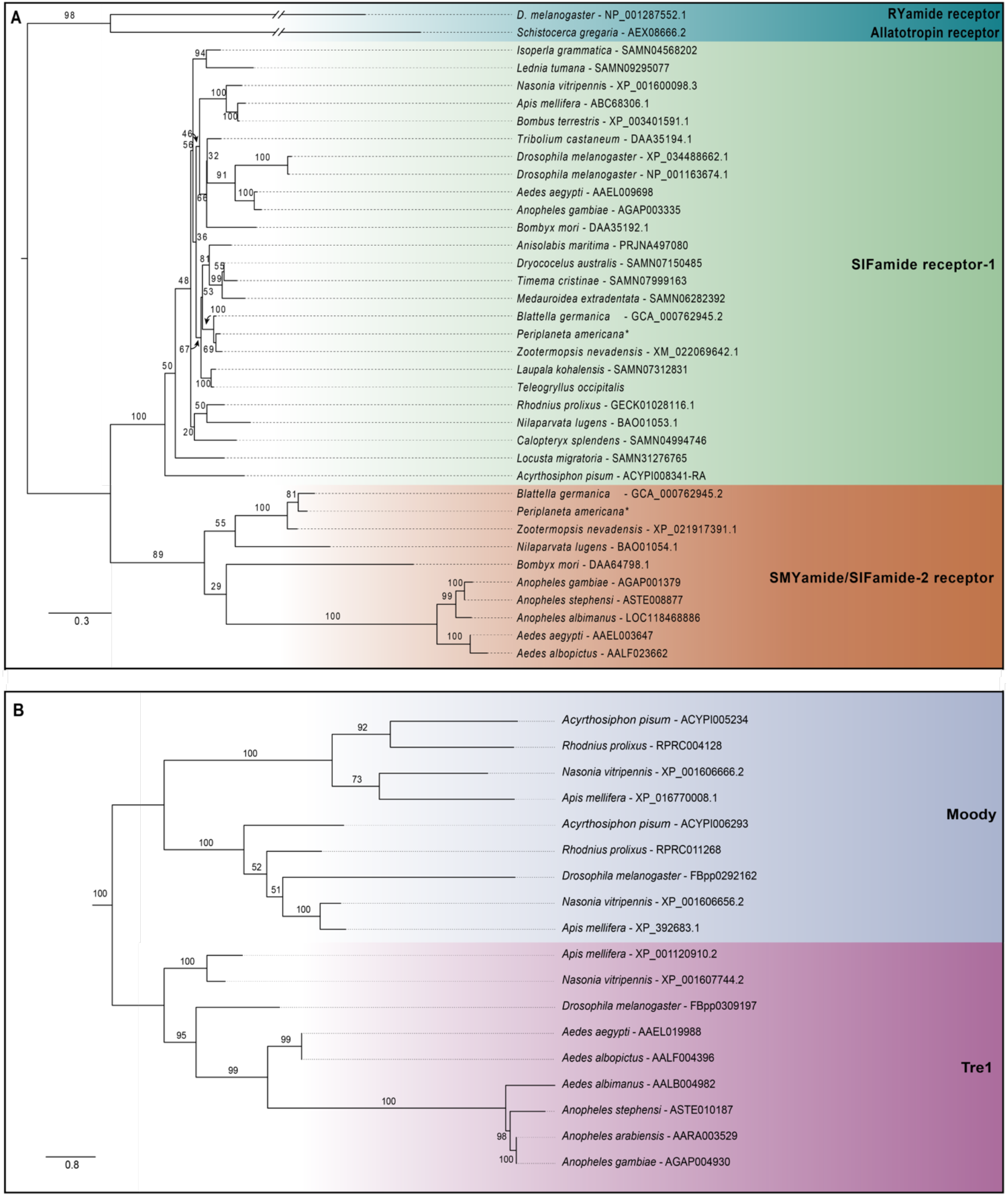
Maximum likelihood tree of **(A)** AAEL003647, **(B)** AAEL019988, and their orthologs. Sequences were aligned using MAFFT version 7.450. Phylogenetic trees were built using PhyML with 100 bootstraps. Bootstrap support is indicated at each node. Sequence accession numbers are provided following the species name of each organism and in table S1. Asterisks after species names indicate sequences that were obtained from Veenstra 2021.

Our analysis identified AAEL019988 as an ortholog of the *D. melanogaster tre1* GPCR with strong support (100/100 bootstraps). *tre1* appears to be highly conserved among holometabolous insects examined in our analysis, including the parasitoid wasp, *Nasonia vetripennis*, the honeybee *Apis mellifera*, and all of the examined dipterans. The genomes of the hemimetabolous insects we examined, *R. prolixus* and *A. pisum*, do not appear to encode an ortholog of *tre1* (figure 1B).

### 3.2 Tissue tropism of orphan receptors

We investigated expression patterns of *AAEL003647* and *AAEL019988* among life stages, sexes, and tissues. Expression of *AAEL003647* was highest in females relative to males and immature stages (one-way ANOVA, *p* < 0.0001) (figure 2A). Expression of *AAEL019988* was higher in adult females relative to 1^st^, 3^rd^, 5^th^ instar larval, and pupal stage mosquitoes (one-way ANOVA, *p* < 0.05). There was no significant difference in expression between females and males (figure 2B). We next examined tissue tropism of the receptors in females. The highest expression of *AAEL003647* and *AAEL019988* was observed in the ovaries (figure 3A-B). We next measured receptor expression across a time series following a blood meal. Our results demonstrate that expression of *AAEL003647* was highest in non-blood fed, 2h, 4h, and 6h pbf female ovaries (figure 3C). Expression of *AAEL019988* was highest in NBF ovaries (figure 3D).

**Figure 2.**
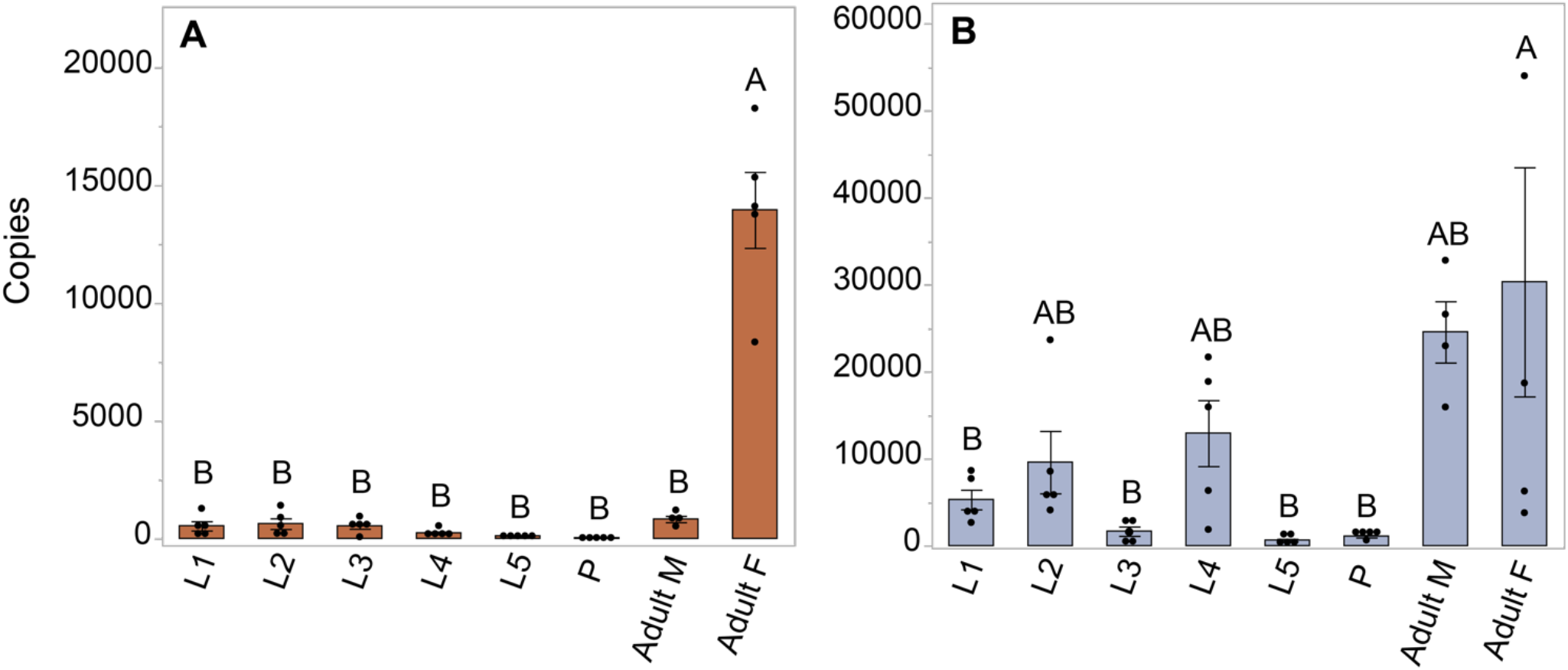
Expression profile of AAEL003647 and AAEL019988 in whole bodies of mosquitoes across life stages and sexes. The x-axis represents the number of copies of AAEL003647 and AAEL019988 per 100ng of RNA. **(A)** Expression of AAEL003647 is significantly higher in adult females (one-way ANOVA, *p* < 0.0001). **(B)** Expression of AAEL019988 was also significantly higher in adult females relative to 1st, 3rd, and 5th stage larvae and pupae (one-way ANOVA, *p* < 0.05). Treatments connected by the same letter are not significantly different (*p* > 0.05, one-way ANOVA).

**Figure 3.**
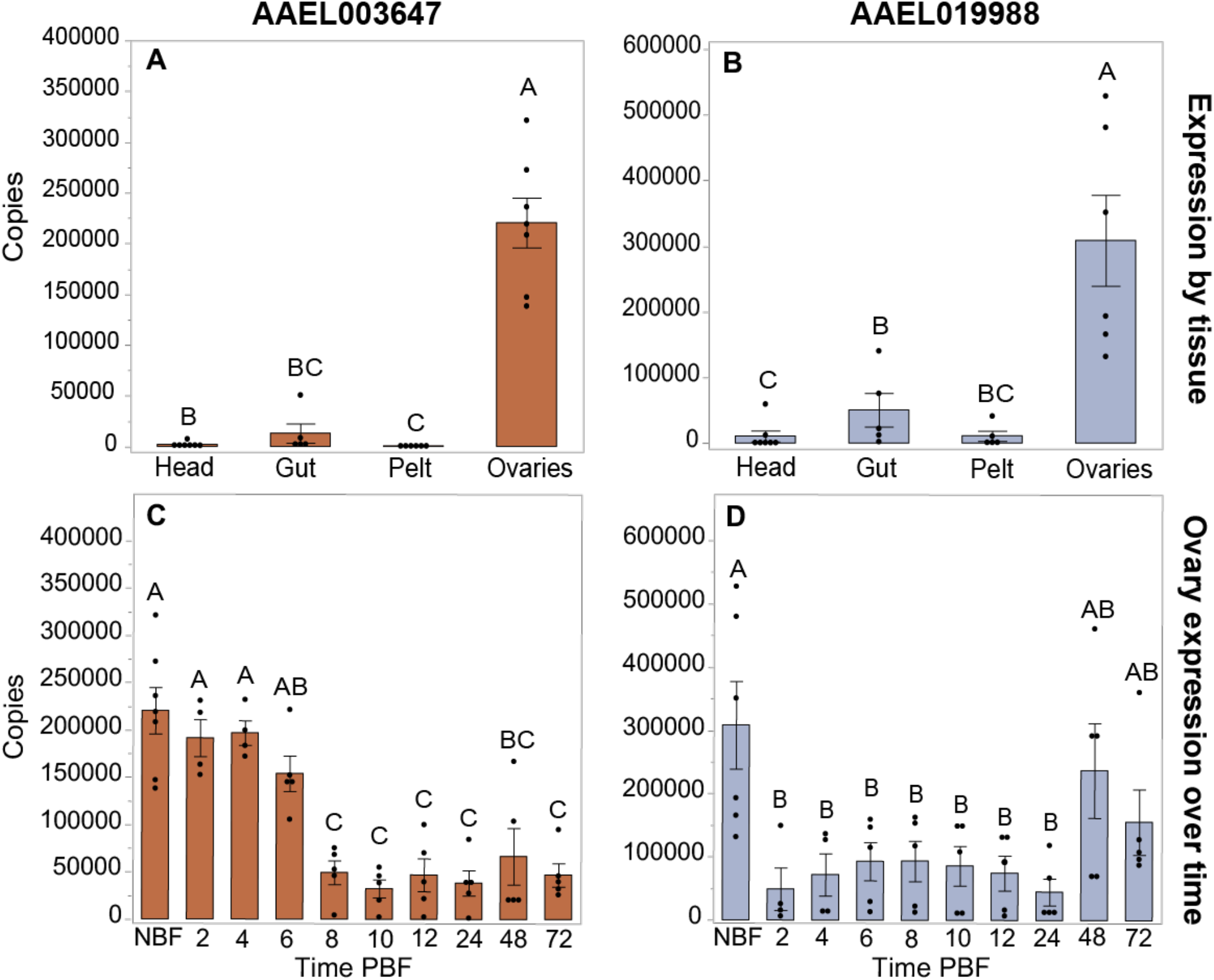
Expression profiles of AAEL003647 and AAEL019988 in NBF *Ae. aegypti* tissues **(A, B)** and in whole bodies following a blood meal **(C, D)**. Expression of AAEL003647 and AAEL019988 is highest in the ovaries for **(A)** AAEL003647 (one-way ANOVA, *p* ≤ 0.003) and **(B)** AAEL019988 (one-way ANOVA, *p* ≤ 0.0092). **(C)** Expression of AAEL003647 is significantly higher in the ovaries of NBF, 2h, 4h, and 6h pbf females (one-way ANOVA, *p* < 0.05). **(D)** Expression of AAEL019988 is significantly higher in the ovaries of NBF females (one-way ANOVA, *p* < 0.05).

### 3.3 Effects of knockdown of AAEL003647 and AAEL019988 on female reproduction

The peaks of expression prior to feeding and nearing the time of oviposition informed our hypothesis that AAEL003647 and AAEL019988 may be important in regulation of egg production and/or oviposition. To understand the effects of both orphan GPCRs on oviposition, we injected newly eclosed female mosquitoes with 2 µg of *ds3647, ds19988*, or *dsEGFP*. For each receptor, we were able to achieve an 85% whole body transcript knockdown (one-way ANOVA, *p* < 0.0163, *p* < 0.0163, respectively; figure 4A-B). Following dsRNA injection, females were allowed to mate and were fed 3 days post-injection. After feeding, females were separated into individual enclosures for oviposition assays. We found that *ds3647* and *ds19988* injected females laid significantly fewer eggs than *dsEGFP* injected females (one-way ANOVA, *p* = 0.0184, *p* = 0.0393, respectively; figure 4C).

**Figure 4.**
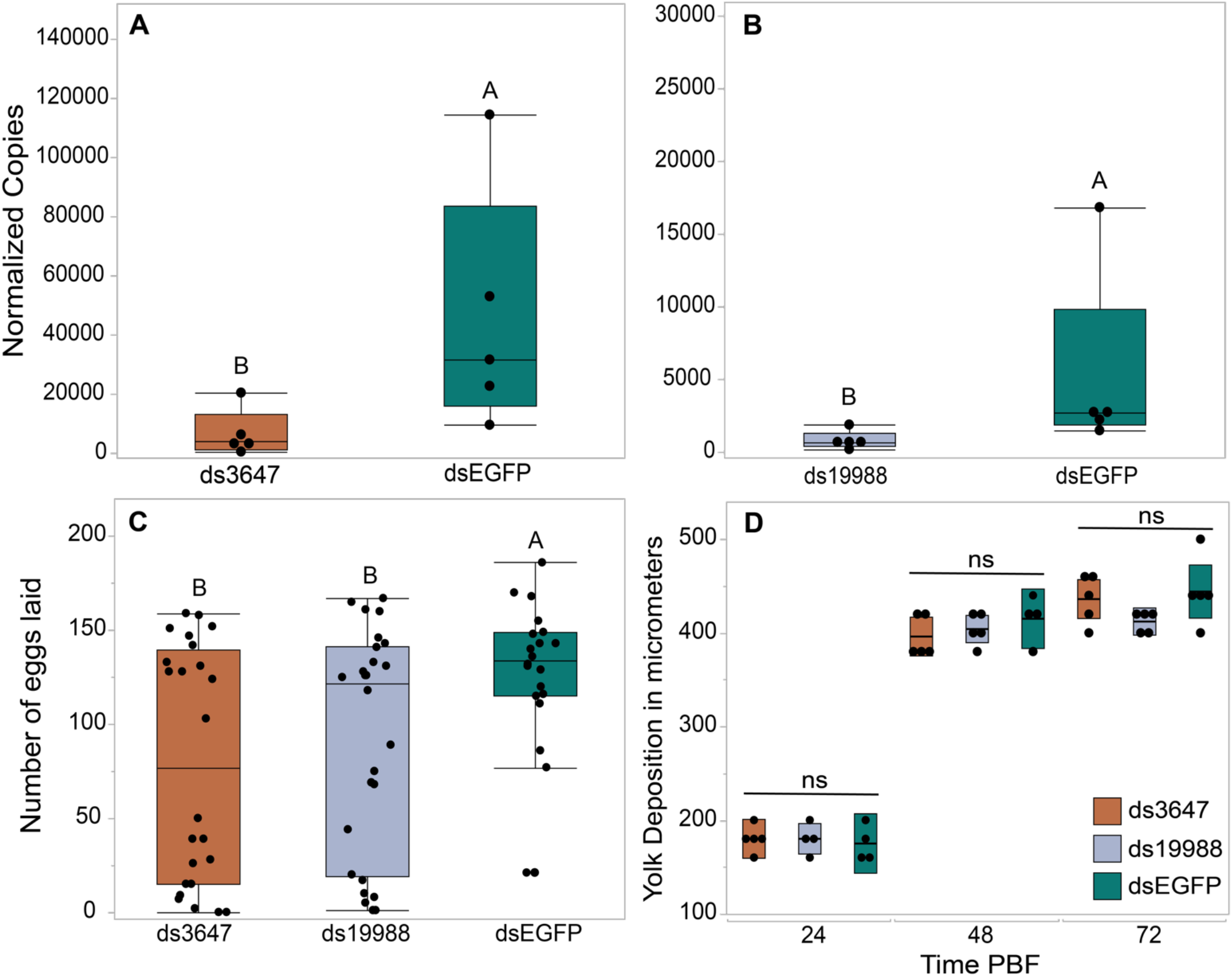
RNAi knockdowns **(A-B)**, oviposition bioassays **(C)**, and yolk deposition **(D). (A-B)** Receptor knockdown validation. We achieved an 85% whole body transcript knockdown for AAEL003647 **(A)** (Wilcoxon rank-sum test, *p* = 0.0163) and AAEL019988 **(B)** (Wilcoxon rank-sum test, *p* = 0.0163). The x-axis represents the number of copies of AAEL003647 and AAEL019988 per 100ng of RNA. Transcripts were normalized by ribosomal S7 expression. **(C)** Knockdown of AAEL003647 and AAEL019988 resulted in a significant decrease in the number of eggs laid relative to dsEGFP controls (Wilcoxon rank-sum test, *p* = 0.0184, *p* = 0.0393, respectively). **(D)** Knockdown of AAEL003647 and AAEL019988 had no effect on yolk uptake (Wilcoxon rank-sum test, *p* > 0.05).

The observed reduction in egg laying by mosquitoes treated with *ds3647* or *ds19988* could be due to a disruption of egg maturation or egg laying. To disentangle this, we examined whether yolk deposition of *ds3647* and *ds19988* injected females was impaired, which would suggest that the receptors are important in post-vitellogenic egg development. We injected newly eclosed females with *dsEGFP, ds3647* or *ds19988*, fed females a blood meal at 3 days post injection, and dissected ovaries from blood fed females at 24, 48, and 72h pbf. Following dissection, we measured the packaged yolk in per individual oocyte with an ocular micrometer. We found no significant difference among oocyte yolk lengths in *ds3647, ds19988*, or *dsEGFP* injected females (one-way ANOVA, *p* > 0.05; figure 4D), suggesting that the receptors mediate physiological events after egg maturation.

## 4. Discussion

Our phylogenetic analysis identified that ancestor of SIFaR underwent gene duplication early in arthropod evolution. This paralog is retained in several arthropod lineages including members of the Culicidae, *Ae. aegypti* (AAEL003647) and *Anopheles gambiae* (AGAP003335). The SIFamide receptor binds the peptide hormone SIFamide, which is localized to neurosecretory cells in the insect brain and central nervous system^29,50^. SIFamide is conserved among hemimetabolous and holometabolous insects and acts as a neurohormone to modulate appetitive behavior^28^, feeding^29^, heart contractions ^29^, and mating behavior^30,51^. The phylogenetic relationships of insect SIFaR receptors indicate an ancient divergence early in arthropod evolution, as evidenced by the presence of two receptor genes in diverse insect species including aphids, cockroaches, and mosquitoes. Veenstra recently identified a novel peptide hormone, SMYamide, in the genome of the American cockroach *Periplaneta americana*^49^. Phylogenetic analysis of the novel peptide revealed that it was sister to the *P. americana* SIFamide peptide, and though binding assays were not performed, the results suggest that SMYamide likely binds the protein encoded by the *SIFaR-2* gene of *P. americana*. Our expanded phylogenetic analysis indicates that the *P. americana* SIFaR-2 is an ortholog of AAEL003647, though we could not identify an ortholog of SMYamide in the *Ae. aegypti* genome. Future binding studies of AAEL003647 will focus on determining if the receptor binds SIFamide, a distant ortholog of SMYamide, or a novel peptide hormone.

The *Drosophila melanogaster* orphan GPCR, Trapped in Endoderm 1 (Tre1), was identified as an ortholog of AAEL019988 in our phylogenetic analysis. Tre1 is essential for the transepithelial migration of germ cells through the posterior midgut during embryogenesis^52–56^. Tre1 is also important for the initiation of courtship behavior *D. melanogaster*^57^. The role of Tre1 in germ cell migration and in courtship may have led to the co-option of this signaling system to regulate reproduction in *Ae. aegypti*. Interestingly, Tre1 is absent in the hemimetabolous insects included in the study, with the closest orthologs identified in their genomes were orthologs of the *D. melanogaster* GPCR *moody*.

Our expression profiles of *AAEL003647* and *AAEL019988* indicate that transcript abundance of both receptors is highest in adult females’ reproductive tissues, suggesting potential roles in egg production. To determine the potential roles of each orphan receptor in female reproductive physiology, we carried out a series of knockdown experiments which resulted in fecundity reduction in *ds3647-* and *ds19988*-injected females. Subsequently, we found that knockdown of both orphan receptors did not affect the amount of yolk packaged into oocytes, suggesting limited interactions with ILP3 and OEH, which are reproductive hormones that are known to modulate oogenesis^1–3^. These results point to a role in oviposition rather than egg production.

The role of the SIFamide, a sister clade to AAEL003647, provides potential clues towards the mechanism of this receptor and its as-yet unknown ligand. SIFamide has been implicated in modulation of feeding and mating behavior in *Drosophila*^28,29,51^. SIFamidergic neurons are activated during starving conditions and are inhibited by the myosin inhibitory peptide (MIP) which modulates satiation^28^. This SIFa/MIP neuropathway governs feeding behavior in *Drosophila*, but also directly affects mating behavior ^28,29,51^. SIFa acts on *fruitless* in *Drosophila*, which modulates courtship behavior; upon inhibition of SIFaR, male flies exhibited bisexual mating behaviors^30,51^. Although AAEL003647 and SIFaR belong to phylogenetically sister clades, it does not guarantee functional similarity. However, there is a possibility these receptors share similar functions, including modulation of oviposition by interaction with MIP.

AAEL019988 is an ortholog of Tre1, which in *Drosophila* regulates mating behavior. Luu *et al*. 2016 found that some *fruitless* expressing neurons also expressed *Tre1*, and that male and female flies exhibited expression of *Tre1* in a sexually dimorphic fashion^57^. Female *Tre*1 expression was induced in males by generating transgenic males expressing the female Tre1 splice form, tra^f^. This *Tre1* “feminization” in males resulted in latency in initiation of courtship behavior and complete absence of courtship initiation behavior in some males. However, there was no significant effect of *Tre1* feminization on the number of offspring per *Tre1* mutant male that mated with a female^57^. We found that knockdown of AAEL019988 disrupts egg laying but not egg development, suggesting that it may have evolved an alternative function not involved in mating behaviors in *Ae. aegypti*. Future studies of AAEL003647 and AAEL019988 will examine the impacts of these orphan receptors on feeding and mating behavior, including through interactions with *fruitless* in *Ae. aegypti*.

## Acknowledgments

The authors would like to express their gratitude to Jena Johnson, Logan Harrell, Lilith South, and Severen Brown for their maintenance of the mosquito colony used for this study.

## Author Contributions

Conceptualization, K.J.V.; methodology, N.I.K.-S. and K.J.V.; experimentation, N.I.K.-S., K.R.S., L.R.A., and K.J.V.; writing—original draft preparation, N.I.K.-S.; writing—review and editing, K.J.V., N.I.K.-S., K.R.S., and L.R.A.; project administration, K.J.V. All authors have read and agreed to the published version of the manuscript.

## Conflicts of Interest

The authors declare that the research was conducted in the absence of any commercial or financial relationships that could be construed as a potential conflict of interest.

## Funding

This research was grant funded by the National Institutes for Allergy and Infectious Diseases of the National Institutes of Health awarded to KJV (Award #K22AI127849).

---

**Table S1:**
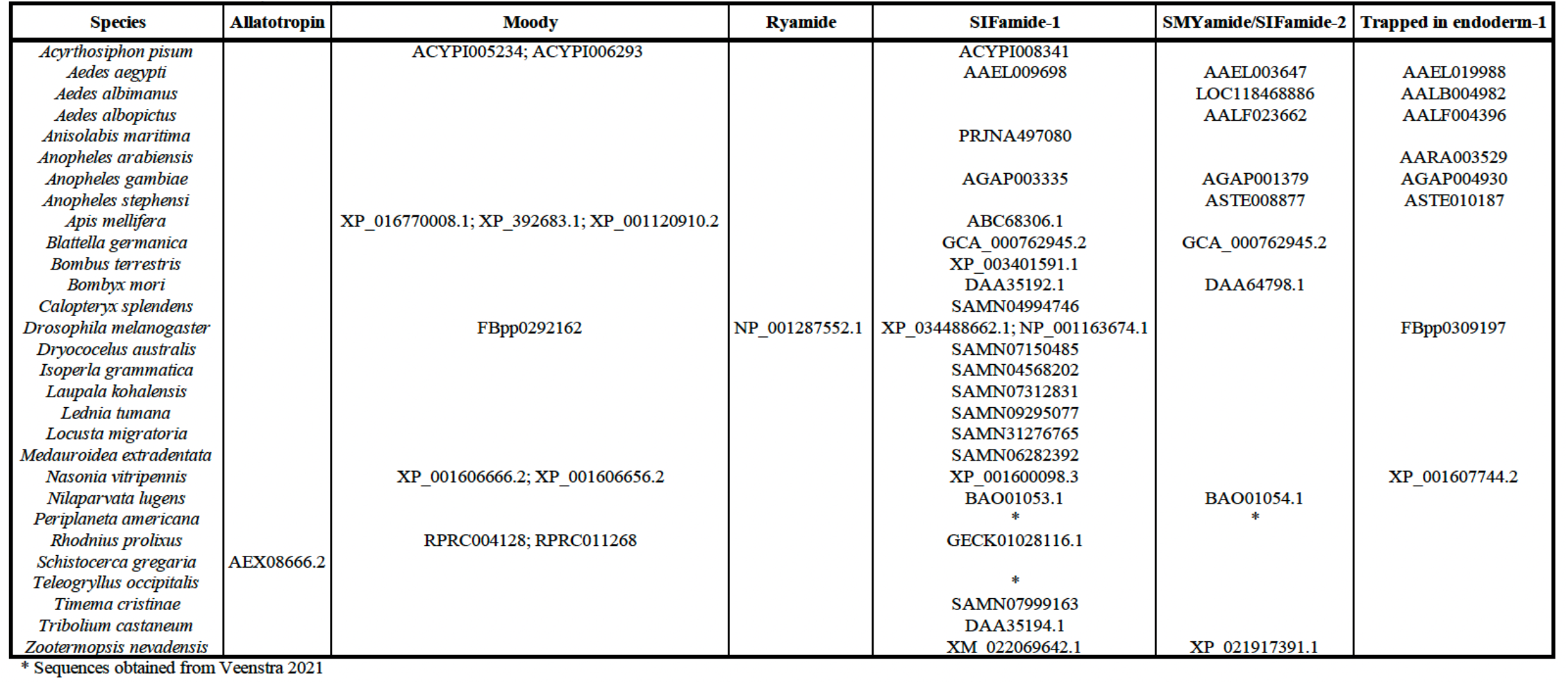
Accessions of protein sequences used in phylogenetic analyses.

**Table S2:**
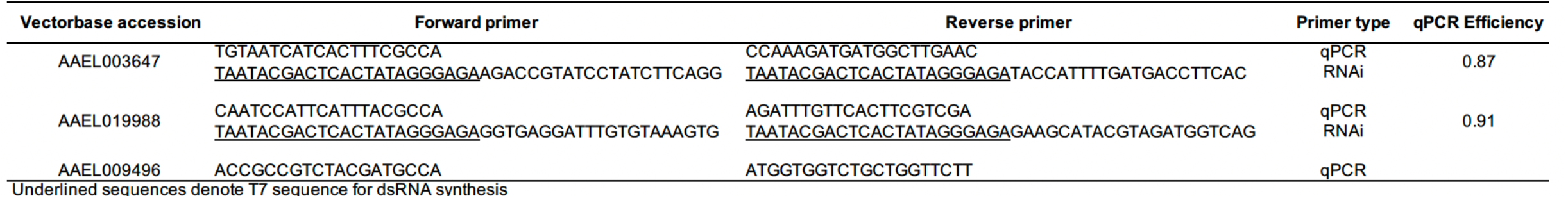
Primers used in this study.

## Notes

### Competing Interest Statement

The authors have declared no competing interest.

